# Human olfactory cortex contributes to emotional and perceptual aspects of aversive associative learning and memory

**DOI:** 10.1101/193748

**Authors:** Yuqi You, Lucas R. Novak, Kevin Clancy, Wen Li

## Abstract

The role of the sensory cortex, beyond the amygdala, has been increasingly recognized in animal associative learning and memory. Here, we examined olfactory cortical plasticity in human olfactory associative learning and memory, while elucidating related changes in emotional and perceptual responses. Psychophysical and neurometric analyses were conducted across an odor-morphing continuum, with the two extreme levels differentially conditioned with aversive and neutral stimuli. Conditioned odors acquired distinct emotional values, tracked by ensemble response patterns in the orbitofrontal (high-level) olfactory cortex. Also observed were enhanced perceptual discrimination and divergent ensemble neuronal response patterns in the anterior and posterior piriform (low-to-intermediate-level) olfactory cortices. Whereas emotional-learning-related changes, both behavioral and neural, maintained 8 days later, perceptual-learning-related changes, also both behavioral and neural, recovered by then, highlighting the human aptitude of forming persistent emotional memory and related sensory cortical plasticity in contrast to transient perceptual alterations of sensory stimuli associated with mild aversive experiences.

## INTRODUCTION

Aversive associative learning via conditioning generates reliable fear/threat learning and memory such that a conditioned stimulus (CS) takes on threat value and prompts conditioned responses of avoidance or attack. Decades of neuroscience research has reached a strong consensus: the amygdala plays a critical role in the acquisition and consolidation of threat learning and initiates and controls conditioned responses (LeDoux, 2000). These findings have generated prominent neural models of anxiety and depression, shedding important light on the pathophysiology of the emotional disorders (Davis, 1992; Dunsmoor and Paz, 2015). However, growing evidence is expanding this dominant view, incorporating the sensory system including the sensory cortex (Grosso et al., 2015a; Li, 2014; Weinberger and Bieszczad, 2011) and, to some extent, the thalamus (Do Monte et al., 2015; Penzo et al., 2015; Weinberger, 2004) as additional neural substrates for aversive associative learning and memory.

Early animal electrophysiology research, dated to the 1950s, demonstrated plasticity in the primary and secondary auditory cortex via associative learning (Diamond and Weinberger, 1984; Galambos et al., 1955; Kraus and Disterhoft, 1982; Weinberger et al., 1984). Recent investigations have rekindled interest in this topic, corroborating and extending the early findings of associative sensory cortical plasticity to all sensory modalities in both humans and animals (Dunsmoor and Paz, 2015; McGann, 2015; Miskovic and Keil, 2012; Ohl and Scheich, 2005; Wilson and Sullivan, 2011). Animal evidence further suggests that this associative plasticity in the sensory cortex not only emerges immediately after conditioning but also shows long-term retention with growing specificity to the CS (Weinberger, 2004). Importantly, associative plasticity in the *secondary* sensory cortex, both immediate and long-lasting, plays a critical role in threat memory, with the immediate plasticity being necessary for the formation (Cambiaghi et al., 2016a; Grosso et al., 2015b; Yang et al., 2016) and the lasting plasticity necessary for the long-term storage (Cambiaghi et al., 2016b; Grosso et al., 2015b, 2017; Kwon et al., 2012; Sacco and Sacchetti, 2010) of threat memory.

In addition to ascribing threat value to the CS, aversive associative learning can also alter basic perceptual processing (e.g., detection and discrimination) of the CS (vs. non-CS), resulting in associative perceptual learning (Chapuis and Wilson, 2012; Li et al., 2008; McGann, 2015; Padmala and Pessoa, 2008; Wilson and Sullivan, 2011). Such associative perceptual learning has also been linked to sensory cortical plasticity, highlighting improved discrimination of the CS that is accompanied by divergent sensory cortical responses to the CS (vs. non-CS; Chapuis and Wilson, 2012; Li et al., 2008) and depends on the integrity of the sensory cortex (Aizenberg and Geffen, 2013). In comparison to typical (non-associative) perceptual learning that often requires prolonged sensory exposure and repeated training (Gibson and Walk, 1994; Gilbert et al., 2001), associative perceptual learning via conditioning possesses an extraordinary advantage of achieving substantial perceptual gain through only a few exposures to the CS (Huang et al., 2007). Furthermore, such associative perceptual learning has the potential to maximize the benefit of threat learning by facilitating defensive response to the CS via enhanced CS detection (Åhs et al., 2013; Padmala and Pessoa, 2008; Parma et al., 2015) while limiting overgeneralized fear response via improved CS discrimination from similar but invalid cues (Aizenberg and Geffen, 2013; Chapuis and Wilson, 2012; Li et al., 2008). Nevertheless, the intricacy between emotional and perceptual aspects of aversive associative learning via conditioning, along with their corresponding forms of sensory cortical plasticity, has not been clearly elucidated. The ultimate ecological benefits of associative learning would depend on their successful transformation into long-term memory. To date, human evidence of associative sensory plasticity has concerned primarily immediate (Li et al., 2008; Miskovic and Keil, 2012; Padmala and Pessoa, 2008) and short-term (∼ 24 hours) effects (Apergis-Schoute et al., 2014), leaving an open question with respect to long-term associative plasticity in the human sensory cortex.

Olfaction represents a unique sensory system, where sensory plasticity via associative learning has been observed across phylogeny and the entire sensory hierarchy, including the first-order receptor neurons (Davis, 2004; Kass et al., 2013; McGann, 2015; Wilson and Sullivan, 2011). Human olfactory aversive conditioning research, though scarce, has revealed remarkable perceptual improvement and olfactory cortical plasticity (Åhs et al., 2013; Li et al., 2008; Parma et al., 2015). Characterized by sensory encoding based on associative, content-addressable memory (Haberly, 2001; Wilson and Sullivan, 2011), the olfactory brain has been adopted as a model system for studying associative memory (Gluck and Granger, 1993; Haberly, 2001). Therefore, interrogation of associative plasticity in the olfactory cortex promises rich insights into emotional and perceptual learning and memory via conditioning. Here, in a human olfactory aversive conditioning paradigm incorporating a constellation of assessments (risk ratings, skin conductance response/SCR, sensory psychophysics, and functional magnetic resonance imaging/fMRI) and importantly, a retention test after an eight-day delay, we examined olfactory emotional and perceptual learning and memory and their respective olfactory cortical plasticity.

As differential conditioning induces specific conditioned responses and promotes perceptual and neural discrimination (Aizenberg and Geffen, 2013; Chen et al., 2011; Ito et al., 2009; Rescorla, 1976), we differentially conditioned two odors (extreme mixtures on an odor-morphing continuum) with an aversive and a neutral UCS to be the threat-relevant CS (CSt) and safety-relevant CS (CSs), respectively (Figure 1A-C). The three intermediate mixtures not paired with the UCS served as control stimuli (i.e., non-conditioned stimuli/nCS), including nCSt (next to the CSt on the morphing continuum), nCSm (the middle nCS), and nCSs (next to the CSs). Psychometric analyses along the morphing continuum would reveal warped psychological distances along perceptual (Figure 1D) and emotional dimensions (Figure 1E) of the odor space, reflecting perceptual and emotional learning, respectively. Multivariate fMRI analyses including multivoxel pattern analysis (MVPA) and representational similarity analysis (RSA) were employed to assay sensory cortical plasticity, along the perceptual and emotional dimensions, across the human olfactory cortical hierarchy (including, from low to high levels, the anterior piriform cortex/APC, posterior piriform cortex/PPC, and olfactory orbitofrontal cortex/OFC_olf_; Gottfried, 2010; Krusemark et al., 2013).

**Figure 1.**
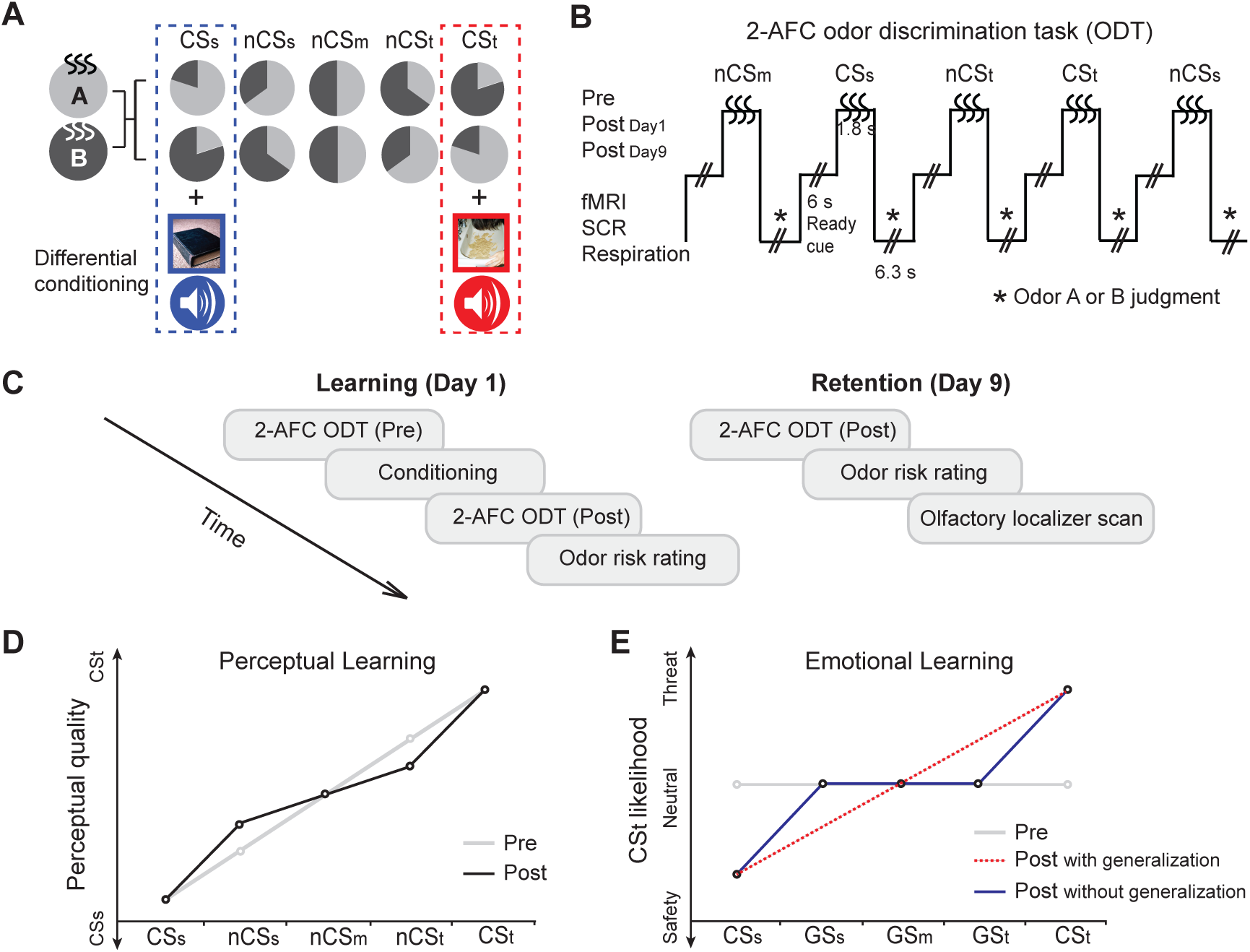
Odor stimuli and experiment design. (A) A continuum of five odor mixtures with systematically morphed proportions (illustrated by the gray wheels) of odors A and B. The two outside mixtures (20% A/80% B and 80% A/20% B) were differentially conditioned as CSt (threat) and CSs (safety) via paired presentation with unconditioned stimuli (UCS: combined sounds and pictures of aversive or neutral emotion), respectively. Assignment of CSt to 20% A/80% B or 80% A/20% B was counterbalanced across subjects. The three intermediate mixtures (35% A/65% B, 50% A/50% B, and 65%A /35% B) served as non-conditioned stimuli (nCS) and were operationalized as nCSt, nCSm, and nCSs based on their distance to CSt or CSs along the odor-morphing continuum. (B) Paradigm of a two-alternative-forced-choice (2-AFC) odor discrimination task (ODT). Fifteen trials of each of five odor mixtures were presented pseudo-randomly for 1.8 seconds, to which participants made judgments of either Odor A or B with button pressing. The task was performed pre-, Day 1 post- and Day 9 post-conditioning, while fMRI, skin conductance response (SCR), and respiration were recorded. (C) Experiment schedule. Day 1 consisted of pre-conditioning 2-AFC ODT, conditioning, post-conditioning 2-AFC ODT, and odor risk rating. Day 9 consisted of post-conditioning 2-AFC ODT, odor risk rating, and an olfactory localizer scan (with a new set of neutral odors). (D) Hypothetical perceptual distances along the odor continuum as warped by conditioning. (E) Hypothetical emotional distances along the odor continuum as warped by conditioning; a tilted slope (red dotted line) or a flat slope (blue solid line) across the three nCS would support conditioning generalization or the lack thereof.

## RESULTS

### Associative perceptual learning and related sensory cortical plasticity

To assess perceptual learning, we administered a two-alternative-forced-choice (2-AFC) odor discrimination task (ODT) before and after conditioning (Figure 1B). Validating the perceptual space along the odor-morphing continuum, a strong linear trend emerged (*F*_1, 31_= 79.62, *P* < .0001), with increasing rates of CSt responses (i.e., endorsing Odor A or B that constituted 80% of the CSt) along the CSs-to-CSt continuum (Figure 2A). To test conditioning-related, associative perceptual learning, we examined discrimination between the CS and their neighboring nCS odors (relative to discrimination between the nCS odors as control conditions to tease out non-associative effects). A psychometric index, the perceptual discrimination index [PDI = (d1+ d4) – (d2 + d3)], was thus derived, where differential CSt response rates (i.e., d’s) between neighboring odors reflected their perceptual distances and the extent of odor discrimination (Figure 2A). Planned contrasts between pre- and post-conditioning PDI revealed a significant increase from pre-to post-conditioning on Day 1 (*t*_31_ = 1.84, *P* = .038, one-tailed; one-tailed tests were applied to all directional hypotheses in the study), but only a weak, nonsignificant trend on Day 9 (vs. the pre-conditioning PDI; *t*_31_ = .94, *P* = .177, one-tailed). These results suggest that conditioning caused short-term perceptual learning, expanding the distance between the CS and their neighboring nCS odors. Notably, despite these distance changes, CSt response rates at Day 1 post-conditioning maintained a strong linear trend along the morphing continuum (*F*_1, 31_= 44.49, *P* < .0001), and considerable perceptual distances remained among the nCS odors: greater CSt response rates for nCSt vs. nCSs (*t*_31_ = 2.05, *P* = .025, one-tailed) and for nCSm vs. nCSs (*t*_31_ = 1.56, *P* = .065, one-tailed). Therefore, conditioning induced quantitative changes in the perceptual space across the odors, leaving their linear configuration largely intact.

**Figure 2.**
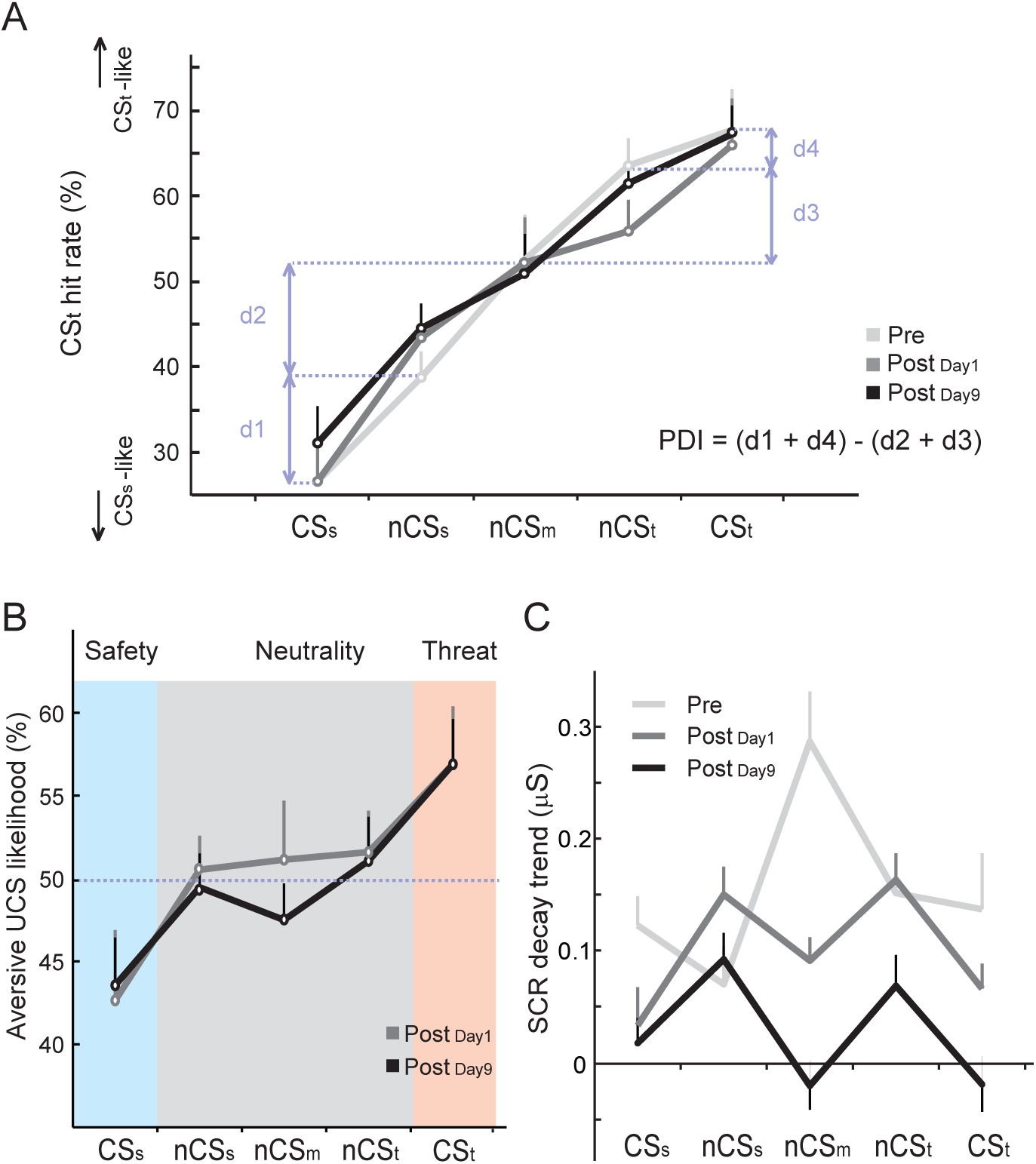
Behavioral and SCR effects of olfactory conditioning. (A) 2-AFC ODT performance at all three time points. The rate of CSt responses (i.e., endorsing Odor A or B that constituted 80% of CSt) at pre-conditioning confirmed a linear increase along the CSs-to-CSt continuum. The perceptual distance (d) between two neighboring odors was reflected by their differential CSt response rate. Perceptual discrimination index (PDI) was calculated by the sum of perceptual distances between the CSt and CSs and their neighbors (d1 + d4) minus the sum of perceptual distances between the nCS and their neighbors (d2 + d3). Conditioning increased PDI from pre-to Day 1 (but not Day 9) post-conditioning. (B) Subjective ratings of the likelihood of receiving aversive UCS following an odor mixture. The patterns at both times post-conditioning conformed to the predicted profile of specific (vs. generalized) conditioning. The three distinct levels, i.e., lowest risk for CSs, chance-level risk for nCS (i.e. nCSs, nCSm, nCSt), and highest risk for CSt, indicated the emergence of distinct affective categories of safety, neutrality, and threat. (C) SCR during the 2-AFC ODT. SCR responses were first fitted with an exponential decay function to account for habituation over repeated trials (Li et al., 2008). Largest SCR decay was observed for nCSm at pre-conditioning, which decreased markedly at post-conditioning (Day 1 and Day 9). Error bars represent s.e.e. (individually adjusted s.e.m.).

To isolate cortical correlates of conditioning-related perceptual learning, we applied RSA to assay dissimilarity (1-Pearson’s *r*) in neural response patterns (i.e., neural distances) to the odors. As illustrated in Figure 3A, we derived an analogous neurometric index, the neural discrimination index [NDI = (d1+ d4) – (d2 + d3); d = 1-*r*], to reflect neural distances between the CS and their neighboring nCS (vs. distances between the nCS as the control condition; Figure 3A). An omnibus ANOVA of Region (APC, PPC and OFC_olf_) and Time (pre-, Day 1 post-and Day 9 post-conditioning) on the NDI revealed a region-by-time interaction (*F*_3.0, 88.7_ = 2.63, *P* = .056; Figure 3B). That is, both APC and PPC exhibited NDI increases from pre- to post-conditioning on Day 1 (APC: *t*_30_ = 1.72, *P* = .048; PPC: *t*_30_ = 1.87, *P* = .035; one-tailed). However, similar to behavioral effects, this effect disappeared on Day 9 (with only a weak, nonsignificant trend in PPC; *t*_30_ = .92, *P* = .184). The OFC_olf_ did not exhibit NDI change at either time point (*P*’s > .293). Therefore, in parallel to perceptual learning, the APC/PPC showed short-term divergence in neuronal population encoding of the CS versus their neighboring nCS.

**Figure 3.**
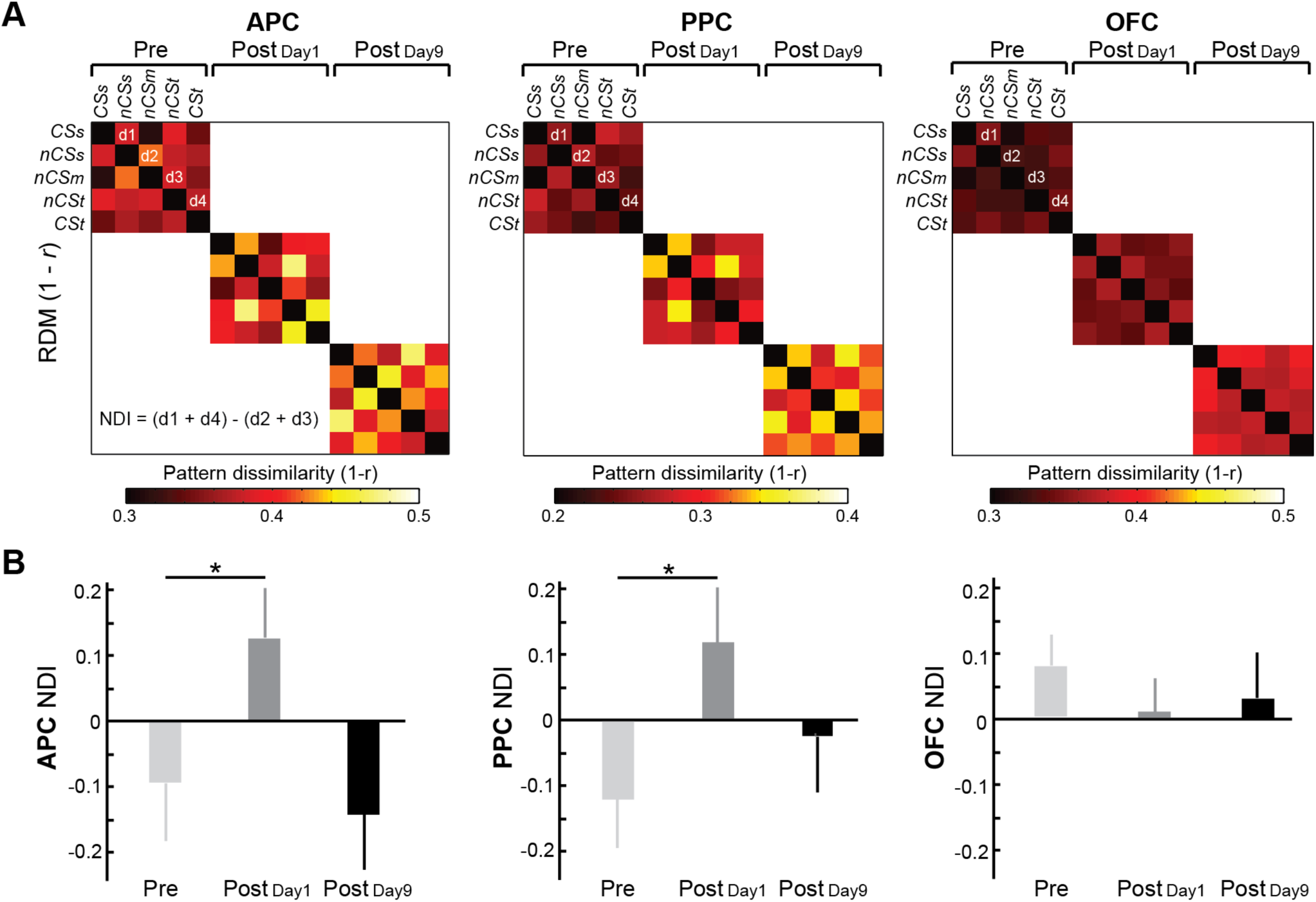
Sensory cortical pattern dissimilarity across the odor space. (A) Group-average representational dissimilarity matrices (RDMs) for APC, PPC, and OFC_olf_ at each time point. Each cell of the matrix reflects pattern dissimilarity (1-Pearson’s *r*) between corresponding odor pairs. Cells right off the diagonal reflect pattern dissimilarity between CSs and nCSs (d1), nCSs and nCSm (d2), nCSm and nCSt (d3), and nCSt and CSt (d4), which were used to calculate a neural discrimination index [NDI = d1 + d4 – (d2 + d3)]. (B) NDI at three time points (pre-, Day 1 post-, and Day 9 post-conditioning) for each ROI. Both APC and PPC demonstrated an increase of NDI from pre- to Day 1 post-, but not Day 9 post-conditioning. Error bars represent s.e.m. * *P* < .05.

### Emotional learning and related sensory cortical plasticity

To assess emotional learning, we acquired post-conditioning risk ratings on a visual analog scale (0-100%) for the five odors, each of which was presented and rated three times, randomly intermixed, to ensure reliability. To minimize response bias and suggestibility, we averted pre-conditioning risk ratings; baseline odor valence ratings confirmed comparable, neutral emotional values for the odors [*P* = .416; valence rating Mean (SD) = 50.6 (19.7)%, on a scale of 1-100%]. An ANOVA (Odor×Time) on risk ratings revealed a main effect of odor (*F*_2.9, 90.5_ = 3.55, *P* = .018; Figure 2B), but no Time or Odor-by-Time effects (*P*’s > .622; ratings on Days 1 and 9 were thus collapsed in the follow-up contrasts). In support of acquired threat and safety meaning, respectively, maximal and minimal risk ratings were observed for CSt and CSs (CSt vs. CSs: *t*_31_ = 3.02, *P* = .005), both differing from chance (50%; CSt: *t*_31_ = 2.67, *P* = .012; CSs: *t*_31_ = −2.40, *P* = .023). Ratings for the three nCS odors were comparable to each other (*F*_1,31_ = .14, *P* = .708), forming a flat slope across them. The ratings hovered around chance (49.3 − 51.3%; *P*’s > .581) and were significantly lower than CSt and higher than CSs ratings (nCS vs. CSt: *t*_31_ = −2.41, *P* = .022; nCS vs. CSs: *t*_31_ = 1.92, *P* = .032; one-tailed). Therefore, akin to the differential conditioning manipulation, these nCS odors showed minimal conditioning generalization but rather merged into an affectively neutral cluster. Together, emotional learning following conditioning resulted in the emergence of three affective classes—threat, safety and neutrality. Critically, the highly similar risk patterns on Day 1 and Day 9 indicate that emotional learning not only arose immediately but also developed into long-term emotional memory.

We also examined SCR responses during the 2-AFC ODT, which, given the task demand of odor discrimination, could index arousal related to difficulty or uncertainty in discriminating the odors. An ANOVA (Odor×Time) revealed main effects of time (*F*_2.0, 59.1_ = 5.09, *P* = .009) and odor (*F*_3.7, 110.3_ = 3.54, *P* = .011) and a time-by-odor interaction (*F*_5.1, 152.7_ = 3.81, *P* = .003). Follow-up ANOVAs for each odor indicated a main effect of time for nCSm (*F*_1.2, 37.4_ = 10.32, *P* = .001) and CSt (*F*_1.8, 54.4_ = 3.24, *P* = .052) but no time effect for the other odors (*P*’s > .155). As illustrated in Figure 2C, the pre-conditioning SCR profile was marked by a maximal response to the nCSm (above all other odors, *P*’s <= .082), in line with the maximal perceptual ambiguity (and hence uncertainty-related arousal) at the midpoint of the morphing continuum. The SCR profile changed drastically post-conditioning on both Day 1 and Day 9, characterized by a large reduction in SCR to nCSm on both Day 1 and Day 9 (*P*’s <= .007). The time effect for CSt was due to a significant SCR reduction from pre- to Day 9 post-conditioning (*P* = .026). Overall, these SCR reductions highlight the largely resolved uncertainty of the nCSm, presumably as this initially ambiguous mid-point odor became distinct being a prototype of the emergent neutral category.

To isolate neural correlates of the three emergent affective categories, we implemented MVPA using the support vector machine (SVM) to classify fMRI response patterns to CSt, CSs and nCSm odors (i.e., category prototypes) during the 2-AFC ODT. Three-class odor classification accuracies were submitted to an omnibus ANOVA (Region×Time), which yielded a Region-by-Time interaction (*F*_3.5, 103.6_ = 3.06, *P* = .025) and was followed with an ANOVA of Time for each region (Figure 4). In support of these emergent categories, classification in the OFC_olf_ improved from the preconditioning baseline, which was at chance [mean (SD) = 32.2 (8.4)%, *P* = .453], becoming reliable at post-conditioning on both days [Day 1/Day 9 mean (SD) = 38.1 (7.4) /36.3 (7.3)%]: Day 1 *t*_30_’s > 3.09, *P*’s < .002; Day 9 *t*_30_’s > 1.99, *P*’s < .031, one-tailed, relative to pre-conditioning or chance levels (Figure 4C, F).

**Figure 4.**
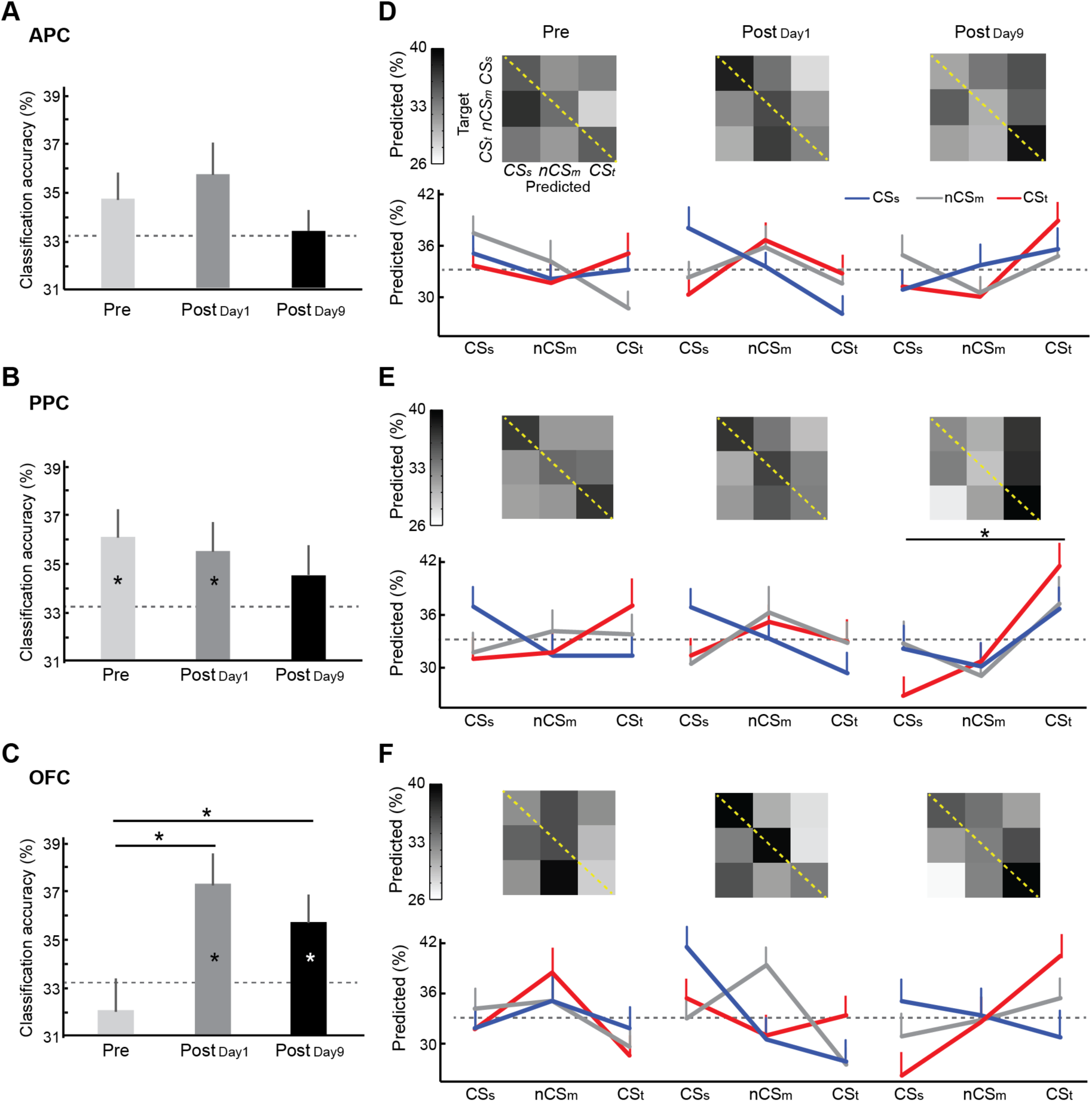
Sensory cortical classification of odors by acquired emotional value. (A-C) Three-class (CSs vs. nCSm vs. CSt) SVM classification accuracies for the APC, PPC, and OFC_olf_ at three time points (pre-, Day 1 post-, and Day 9 post-conditioning). OFC_olf_ showed reliable classification at post-conditioning (both Day 1 and Day 9) but not at pre-conditioning. PPC showed above-chance classification at pre- and Day 1 post-, but not Day 9 post-conditioning. Chance (33.3%) is indicated by a gray dashed line. *: *P* < .05, one-tailed (* inside the bars: vs. chance; * between the bars: post-vs. pre-conditioning). (D-F) Confusion matrices (top) and their line-graph renditions (bottom) for APC, PPC, and OFC_olf_ at three time points. For each confusion matrix, rows represent target (or actual) odors and columns represent the classifier prediction (%) for each odor class (sum = 100%). Diagonal entries (highlighted by yellow dashed lines) indicate correct classification. For line-graph renditions of confusion matrices, each line represents a target odor, with the x-axis indicating the predicted odors. Notably, PPC on Day 9 showed a general bias to classify all odors as the CSt (*: predicted CSt rate > predicted CSs and nCSm rates collapsed; *P* < .05). Chance (33.3%) is indicated by a gray dashed line. Error bars represent s.e.m.

In the PPC, classification was reliable at pre-conditioning [accuracy = 36.1 (6.3)% > chance, *t*_30_ = 2.42, *P* = .011, one-tailed; Figure 4B], confirming the critical role of PPC in odor quality encoding (Gottfried, 2010; Wilson and Sullivan, 2011). At post-conditioning on Day 1, PPC classification [35.5 (6.5)%] remained reliable (> chance: *t*_30_ = 1.85, *P* = .037, one-tailed) and comparable to the baseline (*P* = .749). However, on Day 9, PPC classification reduced to the chance level (*P* = .332), which, as indicated in the confusion matrix (Figure 4E), was caused by an overall bias to classify PPC ensemble responses as CSt-relevant, regardless of the actual odor input (*t*_30_ = 2.74, *P* = .010). Specifically, PPC showed misclassification of CSs and nCSm odors as CSt (vs. the true odors: *t*_30_ = 1.95, *P* = .060) although correct classification for CSt remained high (vs. chance: *t*_30_ = 3.13, *P* = .004). Lastly, the APC did not show reliable classification at pre-conditioning (*P* = .257) or significant change following conditioning (*P*’s > .399; Figure 4A).

### Tuning shifts in the sensory cortex

Stimulus tuning represents a fundamental organization principle in sensory processing, and animal research has revealed neuronal tuning shifts (towards the CS) in the sensory cortex, which consolidates over the course of days to weeks following conditioning (Weinberger, 2004). We thus adopted voxel-based tuning analysis of fMRI data (Serences et al., 2009) to track conditioning-induced tuning shifts.

Based on the pre-conditioning (baseline) response, voxels in a given ROI were sorted into five odor classes by their “optimal odors” (evoking the maximal response among the five odors), after excluding voxels (10%) with lowest mutual information (MI, i.e., non-selective voxels; Serences et al., 2009). A Region-by-Odor ANOVA on the voxel percentages for each class indicated no effect of odor or region, confirming comparable baseline voxel distributions across the odor classes (*P*’s > .202). We then examined two specific voxel classes, tuned to nCSs and nCSt, respectively, concerning their post-conditioning tuning shifts towards the neighboring CS (vs. the neighboring nCS odor—nCSm—as the control; Weinberger, 2004). A tuning shift index (TSI) was thus derived (nCSs TSI = % CSs – % nCSm; nCSt TSI = % CSt – % nCSm) and submitted to a Region-by-Time ANOVA. A Region-by-Time interaction was observed (*F*_1.8, 55.4_ = 4.24, *P* = .022), substantiated by greater TSI in the PPC on Day 9 than Day 1 post-conditioning (*t*_30_ = 2.03, *P* = .026, one-tailed; Figure 5A, B) and no effects of time in the APC or OFC_olf_ (*P*’s > .242). Specifically, the PPC TSI was significant on Day 9 (vs. zero; *t*_30_ = 3.00, *P* = .005) but not on Day 1 post-conditioning (*P* = .802), indicating delayed tuning shifts to the CS. Figure 5C highlights tuning shifts in PPC at post-conditioning Day 9 in a representative participant.

**Figure 5.**
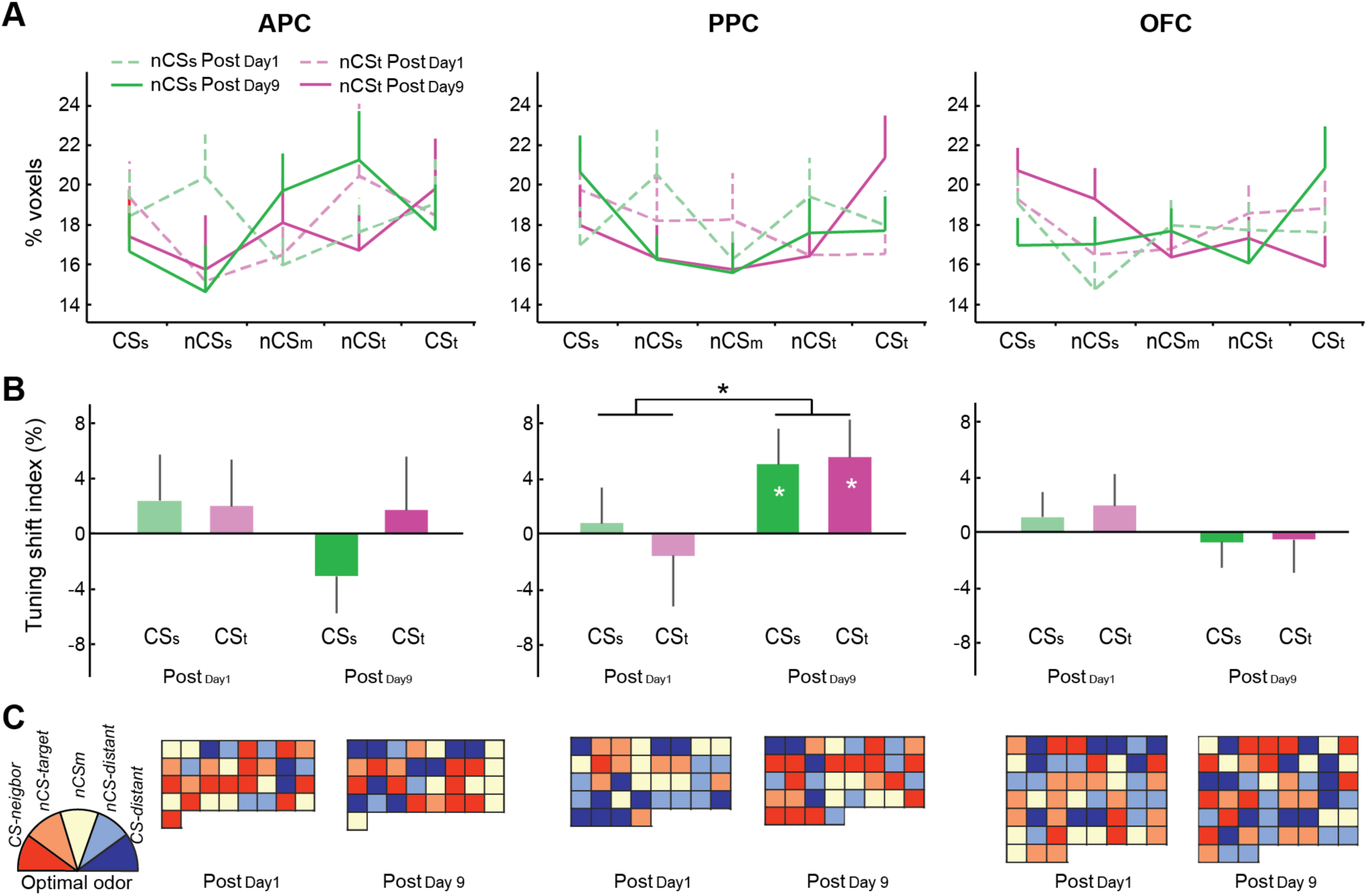
Sensory cortical tuning shifts to the CS. (A) Odor tuning profiles (%) of nCSs and nCSt voxels (defined by the pre-conditioning “optimal odor”) at post-conditioning (Day 1 and Day 9) in the APC, PPC, and OFC_olf_. (B) Tuning shift index (TSI) at post-conditioning (Day 1 and Day 9) for each ROI. TSI scores for the CSs and CSt were defined as (% CSs - % nCSm) and (% CSt - % nCSm), respectively. PPC showed a significant increase in TSI to both CS on Day 9 relative to Day 1 post-conditioning. (C) Odor tuning maps for a representative participant. nCSs and nCSt voxels were combined (denoted as “nCS-target”) and arranged in the descending order of MI (left to right and top to bottom), colors of which indicate their post-conditioning optimal odors. The PPC map at Day 9 post-conditioning contains a disproportional number of red voxels (13/36). Red = “CS-neighbor” (i.e. the adjacent CS: CSt for nCSt, CSs for nCSs); sand = “nCS-target”; light yellow = nCSm; light blue = “nCS-distant” (i.e., the nCS towards the other end of the odor-morphing continuum); dark blue = “CS-distant” (the CS at the other end of the continuum). Error bars represent s.e.m. * *P* < .05, one-tailed (* inside the bars: vs. zero; * between the bars: Day 9 vs. Day 1 post-conditioning).

## DISCUSSION

The growing literature of sensory cortical plasticity via conditioning has expanded our understanding of associative learning, motivating a multi-system, multi-dimension conceptualization of this process. The current study provides some initial insights into this broadened view of associative learning in humans. First, both emotional learning and perceptual learning can arise from aversive conditioning in humans, albeit involving different temporal profiles—immediate and lasting in the former versus immediate but transient in the latter. Second, both threat and perceptual learning recruit the sensory cortex, although engaging dissociable structures in the sensory cortical hierarchy—OFC_olf_ and PPC (delayed involvement) for the former and APC and PPC for the latter. Third, low-level tuning reorganization can develop in the secondary sensory cortex (i.e., the PPC) over time, potentially solidifying sensory cortical representation of the acquired emotional value in the CS.

Differential conditioning promotes divergent conditioned responses to the CS and minimizes conditioning generalization, especially when the CS are related (as in the current study where CSs and CSt shared the same odor components; Aizenberg and Geffen, 2013; Chapuis and Wilson, 2012; Chen et al., 2011). Accordingly, we observe that emotional learning manifested in the threat CS and safety learning in the safety CS, while the intermediate nCS odors exhibited minimal conditioning generalization and merged into a category of emotionally neutral stimuli. Interestingly, the nCSm odor (midpoint of the odor morphing continuum), which evoked a large SCR before conditioning due to the maximal ambiguity in odor discrimination, showed a marked reduction in SCR after conditioning. This effect corroborates emotional learning such that the acquired emotional distinctiveness of the nCSm helped resolve its ambiguity, akin to the idea of “emotion as information” (Damasio et al., 1996). Accompanying this emotional learning, multivoxel patterns of fMRI responses demonstrated that the OFC_olf_, a higher-order olfactory cortex, reliably classified odors containing the acquired affective values (threat, neutrality, and safety) after conditioning. This effect, contrasting the failed classification of these invariant odors at pre-conditioning, indicates associative plasticity in the OFC_olf_ to support emotional learning, consistent with the known function of the OFC in value representation and associative learning (Gottfried and Zald, 2005). Importantly, these convergent multi-modal (risk ratings, SCR, and OFC_olf_ plasticity) effects of emotional learning emerged not only immediately and but also persisted through Day 9, accentuating emotional memory and the contribution of the OFC_olf_ to the storage of acquired emotional value.

The PPC, a secondary (intermediate-level) olfactory cortex, showed reliable odor classification before conditioning, consistent with its critical role in odor quality representation (Gottfried, 2010; Wilson and Sullivan, 2011). While accuracy of PPC odor classification did not change immediately post-conditioning, a striking new pattern of odor classification emerged on Day 9 such that the PPC no longer accurately classified the odors but rather decoded them predominantly as the threat CS odor, regardless of the actual input. While threat and safety values could be acquired via similar mechanisms (Grosso et al., 2015b; Weinberger and Bieszczad, 2011), this classification/decoding bias to threat in the PPC highlights delayed plasticity that may underpin long-term threat memory, potentially facilitating threat memory retrieval by favoring threat decoding or readout. This intriguing disparity in PPC and OFC_olf_ plasticity appears to accord with theories concerning the neural hierarchy of affect, such that intermediate structures (e.g., thalamus) promote complete aversion (to both negative and positive cues) while higher-order forebrain structures maintain balanced affective responses (Smith et al., 2010). In this sense, the olfactory cortical hierarchy may host an affective hierarchy in itself, as the intermediate-level PPC favors threat decoding and promotes defensive behavior while the higher-level OFC_olf_ exerts top-down regulation to optimize emotional responses.

Associative perceptual learning manifested in expanded perceptual distances between the CS and their neighboring nCS odors, as demonstrated in the 2-AFC odor discrimination task. Accompanying this perceptual expansion, dissimilarity (distance) in the multivoxel response patterns increased between the CS and their neighboring nCS odors in the APC and PPC. Both the psychometric and neurometric indices incorporated distances between the nCS odors as controls, thereby largely excluding simple time effects or non-associative perceptual learning via mere sensory exposure. These associative perceptual and neural changes closely resembled previous findings from our lab, where aversive conditioning led to perceptual and neural (PPC) discrimination of the CS odor and a highly similar nCS odor (Li et al., 2008). As APC and PPC represent low and intermediate levels of the olfactory cortical hierarchy, critical for basic encoding of olfactory objects (Gottfried, 2010; Wilson and Sullivan, 2011), their plasticity in parallel to perceptual learning thus highlights updated sensory analysis of olfactory input via associative learning, resulting in altered olfactory perception.

In contrast to emotional learning, this perceptual learning largely preserved the linear perceptual space along the odor-morphing continuum, with the three intermediate nCS remaining somewhat distinguishable perceptual objects. This finding underscores dimensional, quantitative modification of odor perception, accentuating the relative stability of perceptual (vs. emotional) processing. Also in contrast to emotional learning, perceptual learning and related plasticity dissolved over time such that both perceptual and neural spaces across the odor-morphing continuum largely recovered on Day 9, accentuating the constancy of olfactory perception and reliability of basic sensory cortical encoding. Therefore, even though the sensory cortex is sufficiently malleable to exhibit immediate plastic changes to represent acquired biological significance, such changes may succumb to the ecological pressure for the lower-level sensory cortices to maintain fidelity to the external physical world and generate relatively stable percepts. It may require chronic aversive exposures (e.g., repeated conditioning sessions; Parma et al., 2015) or highly traumatic experiences for perceptual learning and related cortical plasticity to develop into stable, long-term memory traces.

Reflecting associative learning on the level of basic sensory processing, stimulus tuning in the PPC shifted to favor the CS odors, consistent with animal data (Weinberger, 2004; Weinberger and Bieszczad, 2011) and a recent human study using scalp electroencephalogram (McTeague et al., 2015). Also coinciding animal findings, this CS-specific odor retuning arose only on Day 9, indicating a time-dependent process likely involved in the transformation of associative learning into long-term memory (Weinberger, 2004). Moreover, the site of this retuning—the PPC, a secondary olfactory cortex—conformed to animal findings, emphasizing conditioning-induced plasticity and long-term memory storage in the secondary sensory cortex (relative to primary or higher sensory levels; Diamond and Weinberger, 1984; Grosso et al., 2015b, 2017; Holschneider et al., 2006; Poremba et al., 1998). The timing (Day 9 only) and site (PPC only) of tuning shifts paralleled changes in higher-order, spatially distributed neuronal ensemble responses in the PPC (i.e., biased PPC classification to the threat CS on Day 9). Synergy between these two processes, threat tuning and preferential threat decoding, could play a critical role in sensory cortical encoding of threat odors in the fashion of content-addressable memory, ensuring sufficient neural and behavioral responses to these threat cues even though the chemical makeup is often diluted, distorted or fragmented by various factors through the journey to the nasal cavity.

In sum, the current study shed some new light on the human associative learning and memory, disentangling its emotional and perceptual aspects and elucidating their corresponding forms of sensory cortical plasticity. Dissociable temporal patterns and cortical substrates are involved in these two aspects of associative learning, serving complementary ecological purposes. Threat learning via conditioning maintains a lasting threat signal to effectively alert the defense system whereas perceptual learning is transient, barring chronic aversive exposures or traumatic experiences (Parma et al., 2015), preserving a reliable cortical code and stable psychological percept for a sensory stimulus. Importantly, the understanding of sensory cortical plasticity via aversive conditioning would promote novel inventions for psychiatric disorders such as post-traumatic stress disorder, by targeting the sensory cortex beyond the limbic-prefrontal-cortical circuitry. The temporal nature of sensory cortical plasticity also highlights a critical period for intervention when the conditioning-related changes are still dynamic and fluid. Timely interventions following trauma could hold special therapeutic promise by preventing early sensory cortical plasticity from consolidating into an enduring content-addressable threat memory.

## MATERIALS AND METHODS

### Participants

Thirty-three individuals (13 males; age 19.9 ± 2.0 years, range 18–25) participated in this two-session fMRI experiment in exchange for course credit or monetary compensation. All participants were right-handed, with normal olfaction and normal or corrected-to-normal vision. Participants were screened to exclude acute nasal infections or allergies affecting olfaction, any history of severe head injury, psychological/neurological disorders or current use of psychotropic medication. All participants provided informed consent to participate in the study, which was approved by the University of Wisconsin-Madison Institutional Review Board. Two participants’ fMRI data were excluded due to metal artefact and excessive movement, leaving 31 participants (11 males) for the fMRI analysis.

### Stimuli

Two generally neutral odorants, acetophenone (5% l/l; diluted in mineral oil) and eugenol (18% l/l) labeled as odors “A” and “B” in the experiment, were systematically mixed to create a morphing continuum of odor mixtures. These two odorants were rated similarly on valence, intensity, familiarity and pungency and have been used as neutral odorants in previous research in our lab (Krusemark and Li, 2013; Novak et al., 2015). The odor continuum consisted of five binary mixtures: 80% A/20% B, 65% A/35% B, 50% A/50% B, 35% A/65% B, and 20% A/80% B (Figure 1A). The two extreme mixtures (20% A/80% B and 80% A/20% B) served as conditioned stimuli (CS), differentially conditioned as CS-threat (CSt) and CS-safety (CSs) via paired presentation with aversive or neutral unconditioned stimuli (UCS; disgusting or neutral images and sounds), respectively. Seven disgusting images (three about dirty toilets and four about human vomit) and seven neutral images (household objects) were selected from the International Affective Picture Set (IAPS; Lang et al., 2008) and internet sources (You and Li, 2016). To strengthen the potency of UCS, each disgust or neutral image was accompanied by a sound of human vomiting or white noise, respectively. Assignment of CSt to either 20% A/80% B or 80% A/20% B was counterbalanced across subjects. The three intermediate mixtures (35% A/65% B, 50% A/50% B, and 65% A/35% B) served as non-conditioned stimuli (nCS), operationalized as nCSt, nCSm, and nCSs based on their distance to the CSt or CSs.

Odor stimuli were delivered at room temperature using an MRI-compatible sixteen-channel computer-controlled olfactometer (airflow set at 1.5 L/min), which permits rapid odor delivery in the absence of tactile, thermal or auditory confounds (Forscher and Li, 2012; Krusemark and Li, 2012; Krusemark et al., 2013; Lorig et al., 1999). Stimulus presentation and collection of responses were controlled using Cogent2000 software (Wellcome Department of Imaging Neuroscience, London, UK) as implemented in Matlab (Mathworks, Natick, MA).

### Two-alternative forced-choice odor discrimination task (2-AFC ODT)

During this task, each odor trial began with a visual get-ready cue, followed by a 3-2-1 countdown, and a sniffing cue, upon which participants were to take a steady and consistent sniff and indicate whether the odor smelled like odor A or B by button pressing (Figure 1B). Each of the five odor mixtures was presented 15 times, in a pseudo-random order with no odor repeated over two trials in succession. Seven additional trials with a central, blank rectangle on the screen (no response required) were randomly intermixed with the odor trials to help minimize olfactory fatigue and establish a non-odor fMRI baseline. Trials recurred with a stimulus onset asynchrony of 14.1 s.

### Experiment procedure

#### Pre-experiment session

Approximately a week prior to Day 1 fMRI session, participants visited the lab. Acetophenone and eugenol were introduced as odors “A” and “B”. All included participants were able to reliably endorse 80% A/20% B and 20% A/80% B as Odors A and B, respectively, after practicing on a 2-AFC ODT with corrective verbal feedback. Participants also provided ratings of the five odor mixtures on valence, intensity, familiarity and pungency using visual analog scales (VAS; 0-10).

#### Experiment Day 1

Participants completed a pre-conditioning 2-AFC ODT, then differential conditioning and a post-conditioning 2-AFC ODT. During differential conditioning, CSt and CSs odors were presented (randomly intermixed, seven trials each) for 1.8 s while the aversive or neutral UCS were presented respectively for 1.5 s at 1s after CS odor onset, with 100% contingency. To prevent extinction, during the post-conditioning 2-AFC ODT (on both Day 1 and Day 9), five extra trials of CSt paired with the aversive UCS were randomly inserted (Li et al., 2008; Onat and Büchel, 2015; Padmala and Pessoa, 2008). Data from these trials were excluded from analysis. After the post-conditioning ODT, each of the five odor mixtures was presented (randomly intermixed, three trials each), to which participants rated on a VAS the likelihood (0-100%) of receiving an aversive UCS following the odor.

#### Experiment Day 9

Participants completed another post-conditioning 2-AFC ODT. After that, another set of UCS risk likelihood ratings was carried out. Finally, we conducted an independent olfactory localizer scan to isolate functional ROIs. Four additional odorants (α-ionone, citronellol, methyl cedryl ketone, 2-methoxy-4-methylphenol), neutral in valence and matched for intensity were presented (15 trials/odor), along with 30 air-only trials pseudo-randomly intermixed, while participants performed a simple odor detection task.

### SCR recording and analysis

During the ODT, skin conductance response (SCR) was continuously acquired at a sampling arte of 1000 Hz using a BioPac MP150 (BIOPAC systems, Goleta, CA) from two MRI-compatible Ag/AgCl electrodes placed on the middle phalanx of the second and third digits of the non-dominant (left) hand. Offline data analysis of SCR waveforms was conducted in Matlab, after low-pass filtering (0.5 Hz) to eliminate MRI scanning artifacts. For each odor trial, evoked SCR response was defined by the magnitude of trough-to-peak SCR deflection during the interval between 0.5 s and 7 s post odor onset, with a minimal evoked deflection of 0.02 µS (Flykt et al., 2007). Since SCR tends to habituate over repeated trials during emotional learning, we modeled the SCR response with an exponential decay function at a rate of ¼ session length (Büchel et al., 1998; Li et al., 2008).

### Respiratory monitoring

Respiration measurements were also acquired (1000 Hz) during the ODT, using a BioPac MP150 with a breathing belt affixed to the participant’s chest to record abdominal or thoracic contraction and expansion. Offline data analysis of respiration waveforms was conducted in Matlab. For each odor trial, a sniff waveform was extracted from a 6 s window post sniff onset and was baseline-corrected by subtracting the mean activity within 1 s preceding sniff onset. Sniff parameters (inspiratory volume, peak amplitude, and peak latency) were generated by averaging across all 15 trials per odor.

### Imaging acquisition and preprocessing

Gradient-echo T2 weighted echoplanar images (EPI) were acquired with blood-oxygen-level-dependent (BOLD) contrast on a 3T GE MR750 MRI scanner, using an eight-channel head coil with sagittal acquisition. Imaging parameters were TR/TE = 2350/20 ms; flip angle = 60°, field of view = 220 mm, slice thickness = 2 mm, gap = 1 mm; in-plane resolution/voxel size = 1.72 × 1.72 mm; matrix size = 128×128. A high-resolution (1×1×1mm^3^) T1-weighted anatomical scan was acquired. Lastly, a field map was acquired with a gradient echo sequence, which was coregistered with EPI images to correct EPI distortions due to susceptibility. Five scan runs, including pre-conditioning, conditioning, Day 1 post-conditioning, Day 9 post-conditioning, and odor localizer, were acquired. Six “dummy” scans from the beginning of each scan run were discarded in order to allow stabilization of longitudinal magnetization. Imaging data were preprocessed in SPM12 (www.fil.ion.ucl.ac.uk/spm), where EPI images were slice-time corrected, realigned, and field-map corrected. Images collected on both Day 1 and Day 9 fMRI sessions were spatially realigned to the first image of the first scan run on Day 1, while the high-resolution T1-weighted scan was co-registered to the averaged EPI of both scan sessions. All multivariate pattern analyses were conducted on EPI data that were neither normalized nor smoothed to preserve signal information at the level of individual voxels, scans, and participants. A general linear model (GLM) was computed on pre-conditioning, conditioning, Day 1 post-conditioning and Day 9 post-conditioning scans. Applying the Least Squares All (LSA) algorithm, we set each odor trial as a separate regressor, convolved with a canonical hemodynamic response function (Abdulrahman and Henson, 2016). Six movement-related regressors (derived from spatial realignment) were included to regress out motion-related variance. For the odor localizer scan, we applied a GLM with odor and no odor conditions as regressors, convolved with a canonical hemodynamic response function and the temporal and dispersion derivatives, besides the six motion regressors of no interest. A high-pass filter (cut-off, 128 s) was applied to remove low-frequency drifts and an autoregressive model (AR1) was applied to account for temporal nonsphericity.

### ROI definition

To delineate learning-induced sensory cortical plasticity, we focused on a set of *a priori* ROIs across the olfactory cortical hierarchy: anterior piriform cortex (APC), posterior piriform cortex (PPC), and olfactory orbitofrontal cortex (OFC_olf_). Amygdala and hippocampus, critical limbic areas involved in acquisition of emotional and contextual associative memory, were included in the supplemental analysis (Figure S1). All five ROIs were manually drawn on each participant’s T1 image in MRIcro (Figure S2; Rorden and Brett, 2000), with the OFC_olf_ in reference to a meta-analysis (Gottfried and Zald, 2005) and prior literature (Howard et al., 2009) and the other ROIs in reference to a human brain atlas (Mai et al., 1997). Left and right hemisphere counterparts were merged into a single ROI. We then applied functional constraints to these anatomical ROIs based on the odor-no-odor contrast conducted on the independent odor localizer scan for each participant, with a liberal threshold at *P* < 0.5 uncorrected (Li et al., 2008).

### Multivariate fMRI analysis

To delineate neural plasticity, we applied multivariate fMRI analysis to characterize changes in ensemble neuronal response patterns to the odor mixtures following conditioning. Two types of multivariate analyses were conducted across the olfactory cortical hierarchy (including, from low to high, the APC, PPC, and OFC_olf_) and over limbic regions (i.e., amygdala and hippocampus). While the multivoxel pattern analysis (MVPA) uses pattern classification techniques to decode the category of objects or distinct psychological states, the representation similarity analysis (RSA) uses correlations across multivoxel patterns of responses to assess the degree of similarity/relation in patterns evoked by different stimuli (Kriegeskorte et al., 2008; Poldrack and Farah, 2015). Therefore, we used the MVPA to isolate the neural underpinnings of the emergent affective categories, and the RSA to identify the neural plasticity that matched the warped psychological space of the odor mixtures following conditioning.

#### Multivoxel Pattern Analysis (MVPA)

We implemented a linear Support Vector Machine (SVM) to perform supervised three-class pattern classification between CSs, nCSm, and CSt. For each ODT in each participant, beta values for each CSs, nCSm, and CSt trial (15 trials per odor, 45 in total) were extracted for all voxels within each functional ROI, and were normalized (Z-scored) across trials for each voxel. Applying leave-one-out cross-validation, we iteratively left out a set of pattern vectors (3 trials/odor) as the testing set and trained a linear SVM (LIBSVM; http://www.csie.ntu.edu.tw/cjlin/libsvm) on the remaining vectors (12 trials/odor). We used the optimal C parameter (determined by an extensive grid-search—2^-11^ ∼ 2^15^) for the SVM, which then classified the three odors based on the testing set. The classification outcome, averaged across 15 iterations, yielded a 3 X 3 confusion matrix. Each row represented a target (or actual) odor, and each column represented the probability of the odor being classified as one of the three possible odors. Accordingly, the diagonal entries indicated correct classifications (i.e., hits) whereas the off-diagonal values indexed mis-classifications (i.e., misses). The overall classification accuracy was defined as the average classification accuracy across the diagonal values.

#### Representational similarity analysis (RSA)

For each ODT in each participant, trial-wise beta values were extracted for all voxels within a functionally constrained ROI. These beta values were averaged across all 15 trials of each odor mixture, resulting in an odor-specific linear vector of beta values across a given ROI. Pearson’s correlation (*r*) was computed between all pairs of pattern vectors (5 odors×3 times), resulting in a 15×15 correlation matrix—the representational similarity matrix. To directly represent neural distances, this matrix was converted into a representational dissimilarity matrix (RDM) by replacing the *r* values with dissimilarity scores (1 – *r*; Fisher transformed). Dissimilarity scores (d’s) between CSs and nCSs (d1), nCSs and nCSm (d2), nCSm and nCSt (d3), and nCSt and CSt (d4) were then extracted to form a neural discrimination index [NDI = d1+d4 – (d2+d3)].

### Tuning analysis

Based on animal tuning analyses (Bakin and Weinberger, 1990; Weinberger et al., 1993), we adapted a voxel-based tuning analysis used for visual sensory encoding (Dubois et al., 2015; Serences et al., 2009) to assess odor tuning in the olfactory sensory cortices. For each ODT in each participant, trial-wise beta values (5 odors×15 trials), after removing the mean beta for a given trial across all voxels in an ROI, were normalized (by z-scoring) across trials. We computed the mutual information (MI) to quantify the amount of information each voxel conveyed about the odors. First, we converted the beta values into a discrete variable (*B*) by dividing the range of betas into a set of equidistant bins (*b*). The size of the bins was determined by Freedman-Diaconis’ rule [bin size = (max(*B*) – min(*B*))/2*IQR*n^-1^/^3^], where n is the number of trials (n = 75). We selected the median bin size of all voxels within an ROI based on pre-conditioning data, and held it constant for the post-conditioning sessions (Day 1 and Day 9). Next, we computed for each voxel the entropy of (discretized) responses (*B*) as follows:

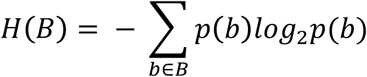

where *p(b)* proportion of trials whose responses fall into bin *b*. Then, we computed conditional entropy, the entropy *H(B\o)*, of responses given knowledge of the odor condition, as follows:

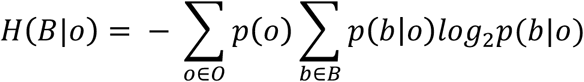

where *p(b\o*) is the proportion of trials falling into bin *b* when responding to a certain odor (*o*). The index of mutual information *M1(B;O)* amount of information the responses of a voxel carries for an odor, was calculated as the reduction in entropy of responses given knowledge of the odor condition:

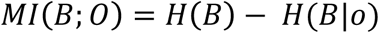

As odor-informative voxels yield high MI, voxels with low MI values suggest no mutual dependence between the distribution of responses (*B*) and odor (*O*), indicating indiscriminant or random responses to all odors. Voxels with bottom 10% MI values in a given ROI were thus excluded as non-informative voxels. For each remaining voxel, its preferred odor (tuning) was defined as the odor eliciting the largest beta among the five odor mixtures. As such, we sorted all the informative voxels at pre-conditioning into five classes. In parallel to animal research (Bakin and Weinberger, 1990), we targeted the two classes of voxels that were tuned to nCSs and nCSt, respectively, before conditioning, and tested whether these voxels shifted their tuning preferences to CS after conditioning. At Day 1 and Day 9 post-conditioning, respectively, we examined percentages (%) of these voxels tuned to each of the five odor mixtures (after excluding the bottom 10% MI, non-informative voxels). If tuning indeed shifted to the CS, we would expect greater % of these voxels tuned to CSs (for nCSs) and CSt (for nCSt), respectively, relative to the neighboring odor, nCSm. As such, we computed a tuning shift index (TSI) for the nCSs and nCSt voxels: % CSs – % nCSm and % CSt – % nCSm, respectively, for Day 1 and Day 9 post-conditioning in each ROI and participant.

## Statistics

Given the considerable sample size, we performed parametric tests here. When multiple factors were involved (e.g., Region and Time) in testing a hypothesis, we would protect the tests with an omnibus ANOVAs (with Greenhouse-Geisser correction for non-sphericity). Specifically, we performed a 2-way ANOVA (Time×Odor) on the SCR and risk ratings, 2-way ANOVAs (Time ×Region) on the NDI and SVM accuracy, and a 3-way ANOVA (Time×Region×Valence) on the TSI scores. Paired *t*-tests were conducted following significant *F* statistics. Planned *t*-contrasts were applied on PDI given the simplicity of these tests. Two-tailed *P*’s were applied to all tests except for those addressing clearly directional hypotheses (i.e., PDI, NDI, SVM accuracy, risk ratings, and TSI).

## AKNOWLEDGEMENTS

This research was supported by the National Institute of Mental Health (R01MH093413 to W.L.). The authors declare no competing financial interests or potential conflicts of interest.

